# Management, Analyses, and Distribution of the MaizeCODE Data on the Cloud

**DOI:** 10.1101/852269

**Authors:** Liya Wang, Zhenyuan Lu, Melissa delaBastide, Peter Van Buren, Xiaofei Wang, Cornel Ghiban, Michael Regulski, Jorg Drenkow, Xiaosa Xu, Carlos Ortiz-Ramirez, Cristina Fernandez-Marco, Sara Goodwin, Alexander Dobin, Kenneth D. Birnbaum, David P. Jackson, Robert A. Martienssen, William R. McCombie, David A. Micklos, Michael C. Schatz, Doreen H. Ware, Thomas R. Gingeras

**Affiliations:** Cold Spring Harbor Laboratory, Cold Spring Harbor, NY; New York University, New York, NY; Johns Hopkins University, Baltimore, MD; USDA-ARS Robert W. Holley Center for Agriculture and Health, Ithaca, NY, United States

**Keywords:** bioinformatics, functional annotations, cloud computing, ENCODE, workflows

## Abstract

MaizeCODE is a project aimed at identifying and analyzing functional elements in the maize genome. In its initial phase, MaizeCODE assayed up to five tissues from four maize strains (B73, NC350, W22, TIL11) by RNA-Seq, Chip-Seq, RAMPAGE, and small RNA sequencing. To facilitate reproducible science and provide both human and machine access to the MaizeCODE data, we enhanced SciApps, a cloud-based portal, for analysis and distribution of both raw data and analysis results. Based on the SciApps workflow platform, we generated new components to support the complete cycle of MaizeCODE data management. These include publicly accessible scientific workflows for the reproducible and shareable analysis of various functional data, a RESTful API for batch processing and distribution of data and metadata, a searchable data page that lists each MaizeCODE experiment as a reproducible workflow, and integrated JBrowse genome browser tracks linked with workflows and metadata. The SciApps portal is a flexible platform that allows the integration of new analysis tools, workflows, and genomic data from multiple projects. Through metadata and a ready-to-compute cloud-based platform, the portal experience improves access to the MaizeCODE data and facilitates its analysis.

## INTRODUCTION

Maize is one of the most biologically, socially, and economically important crop plants. Following the sequencing of its genome (Jiao et al., 2017; Schnable et al., 2009), the next critical step in understanding maize biology will involve identifying and deciphering functional sequence regions. Modeled on the Encyclopedia of DNA Elements (ENCODE) Project for the human genome (ENCODE Consortium, 2004), the MaizeCODE project is an integrated and multi-disciplinary project aimed at revealing the functional regions of the maize genome by identifying loci that are transcribed, methylated, or bound by specific modified histones and transcription factors in various tissues. In addition, MaizeCODE is designed to store, collate, display, and disseminate the data to the wider community of plant biologists worldwide.

To curate, process, and distribute the ENCODE data, the ENCODE Data Coordination Center (DCC) group established the ENCODE portal (Sloan et al., 2016), which relies on both rich metadata and commercial cloud resources through the DNAnexus platform (https://www.dnanexus.com). Within this platform, standard processing pipelines for human genome analysis are constructed to ensure consistent and reproducible processing of primary sequence data. However, both the ENCODE DCC and end users are required to cover the cost of the DNAnexus service and commercial cloud resources. In order to provide a cost-free data processing platform for academic users, the MaizeCODE DCC group decided to leverage two NSF-funded resources, XSEDE (Towns et al., 2014) at the Texas Advanced Computing Center (TACC) for computing power and the CyVerse Data Store (Goff et al., 2011) for cloud-based data storage.

To automate bioinformatics analysis over both the XSEDE/TACC cloud and CyVerse Data Store, we developed a bioinformatics workflow platform called SciApps (Wang et al., 2018). In the work described here, we further improved SciApps by adding a RESTful API for automating batch processing of the MaizeCODE data and metadata management, a searchable MaizeCODE data page powered by a relational database, several analysis workflows, and Genome Browser tracks automatically generated from unique workflow identifiers via the RESTful API. The SciApps platform (https://sciapps.org) has been used to support both MaizeCODE DCC and end users to process/reprocess and manage multi-omics data through either the GUI or the API.

## METHODS

### Overview of the entire MaizeCODE data management cycle

To improve accessibility, reproducibility, reusability, and interoperability, data generated by the MaizeCODE Consortium members are uploaded to a cloud-based data storage system, the CyVerse Data Store (Goff et al., 2011). The Data Store, which is built on top of iRODS (a rule-oriented data system) (Moore and Rajsekar 2010), supports data virtualization, sharing, bulk uploads/downloads, and collaborations. Once uploaded, the experimental metadata are attached to raw data files to facilitate the reuse of data, as well as submission of data to the NCBI Short Read Archive (SRA). The metadata is later retrieved via the Terrain API (https://github.com/cyverse-de/terrain) to automate batch analyses via SciApps (Wang et al., 2018). SciApps provides a ready-to-compute cloud-based platform for automating complex analyses constructed using modular applications (or apps). Previously, SciApps was operated via a web GUI, but since that time we have developed SciApps RESTful APIs to support batch processing of MaizeCODE and other data. In addition, we have integrated over 20 new apps for ground-level analyses such as quality control (QC), alignment to the reference genome, filtering, quantification (e.g. for gene expression), and peak calling (if needed). Replicates (and controls if available) of each assay are organized as a single experiment (or workflow with a unique ID), which represents an entity that chains together raw data, analysis results, experimental metadata (such as tissue, assay, etc), and computational provenance (computational metadata supporting the reliable replication of scientific results, such as version of the software tools, parameters used for the analysis, etc). SciApps extracts experimental metadata and attaches them to a specific workflow so that users can access them directly on the SciApps portal. Genome browser tracks are automatically generated and displayed within an integrated version of JBrowse (Skinner et al., 2009) by looping through the list of experiments/workflows via the SciApps RESTful API.

In summary, the SciApps portal and RESTful API have been used to support the management, analysis, and distribution of the MaizeCODE data. Through the automation of both data and metadata management, the chances of human errors in data management are greatly reduced.

### Processing and accessing the MaizeCODE data with the SciApps RESTful API

The cloud-based architecture of SciApps (Wang et al., 2018) enables highly scalable processing of MaizeCODE data on the XSEDE/TACC cloud. Both intermediate and final results are archived in the CyVerse Data Store, where the raw data are also hosted. As discussed above, each SciApps workflow captures experimental metadata and computational provenance along with the raw and processed data. Batch processing of MaizeCODE data is supported through the RESTful API via a workflow endpoint that takes a template workflow (for example commands, see Supplementary file page 3); the API endpoints are provided in **Table 1**.

**Table 1.** SciApps release 1.0 RESTful API

| Endpoint | Method | Description |
| --- | --- | --- |
| /job | GET | List all jobs |
| /job/new/{id} | POST | Run a new job |
| /workflow/build | POST | Build a workflow from jobs |
| /workflowJob/new | POST | Generate a workflow JSON |
| /workflow/new | POST | Save a new workflow |
| /job/{id} | GET | Return the job JSON |
| /job/{id}/delete | GET | Delete the job |
| /workflowJob/run/{id} | GET | Run a new workflow |
| /workflow/{id}/metadata | GET | Get the workflow metadata |
| /workflow/{id}/update | POST | Update the workflow with metadata etc |
| /workflow | GET | List all workflows |
| /apps/{id} | GET | Return the application JSON |
| /workflow/{id}/delete | GET | Delete the workflow |
| /workflow/{id} | GET | Return the workflow JSON |
| /apps | GET | List all integrated apps |

The analysis workflow for a specific assay is typically built interactively within the SciApps GUI using one data set as a template. Once the workflow is captured, it can then be easily and automatically applied to analyze other genomes and tissues. Alternatively, users may also build workflows entirely programmatically with a series of analysis job IDs via the API. Experimental metadata are retrieved via the CyVerse Terrain API, and then attached to the workflow via the SciApps API at runtime. The API supports the MaizeCODE DCC for automatically processing a large amount of data and also supports retrieval of results and metadata by end users. For example, genome browser tracks can be automatically generated given a workflow ID by the following steps (**Figure S1**): 1. Retrieve job IDs and inputs with the workflow endpoint, given a workflow ID; 2. Retrieve the output path with the job endpoint, given a job ID; 3. Construct the browser-ready link with the retrieved information. To simplify the process, the MaizeCODE DCC encodes the genome, tissue, and replicate information into the input raw data file path, which is also accessible through the workflow metadata endpoints. SciApps also names the output filename based on the input filename with the output ID (defined by the app) as the prefix. As shown in **Figure S1**, once the input filename, input path, and output path to cloud storage are retrieved by calling the API, the output file path can be constructed to build the browser links.

Given a workflow ID, users can also call the API to retrieve the computational metadata (e.g., https://sciapps.org/workflow/a14ff622-7af9-4b1f-877a-2be926dc1059) or the experimental metadata (e.g., https://sciapps.org/workflow/a14ff622-7af9-4b1f-877a-2be926dc1059/metadata) in standard JSON format and view them in any web browsers.

### Accessing the MaizeCODE experiments as reproducible workflows

The MaizeCODE data page can be accessed under ‘Data’ from the top navigation bar of SciApps (**Figure 1**). Keyword search is supported to allow the user to narrow down the list of experiments to a specific genome or tissue or assay in real time. Once an experiment is selected, the user can access the metadata, workflows, and ground-level analysis results of the experiments, starting from raw sequence data. With the ‘Relaunch’ tab, user can reproduce the entire analysis with one click or apply the same analysis workflow to new data. Using the ‘Share’ tab, the analysis can be shared with others. Users can load the results to the History panel and subject them to further analysis using the modular apps. Because all results are archived in the cloud, downstream analyses can be completed quickly, e.g., differential expression analysis between two tissues can be completed in a few minutes, rather than hours when starting from the raw sequence data.

**Figure 1.**
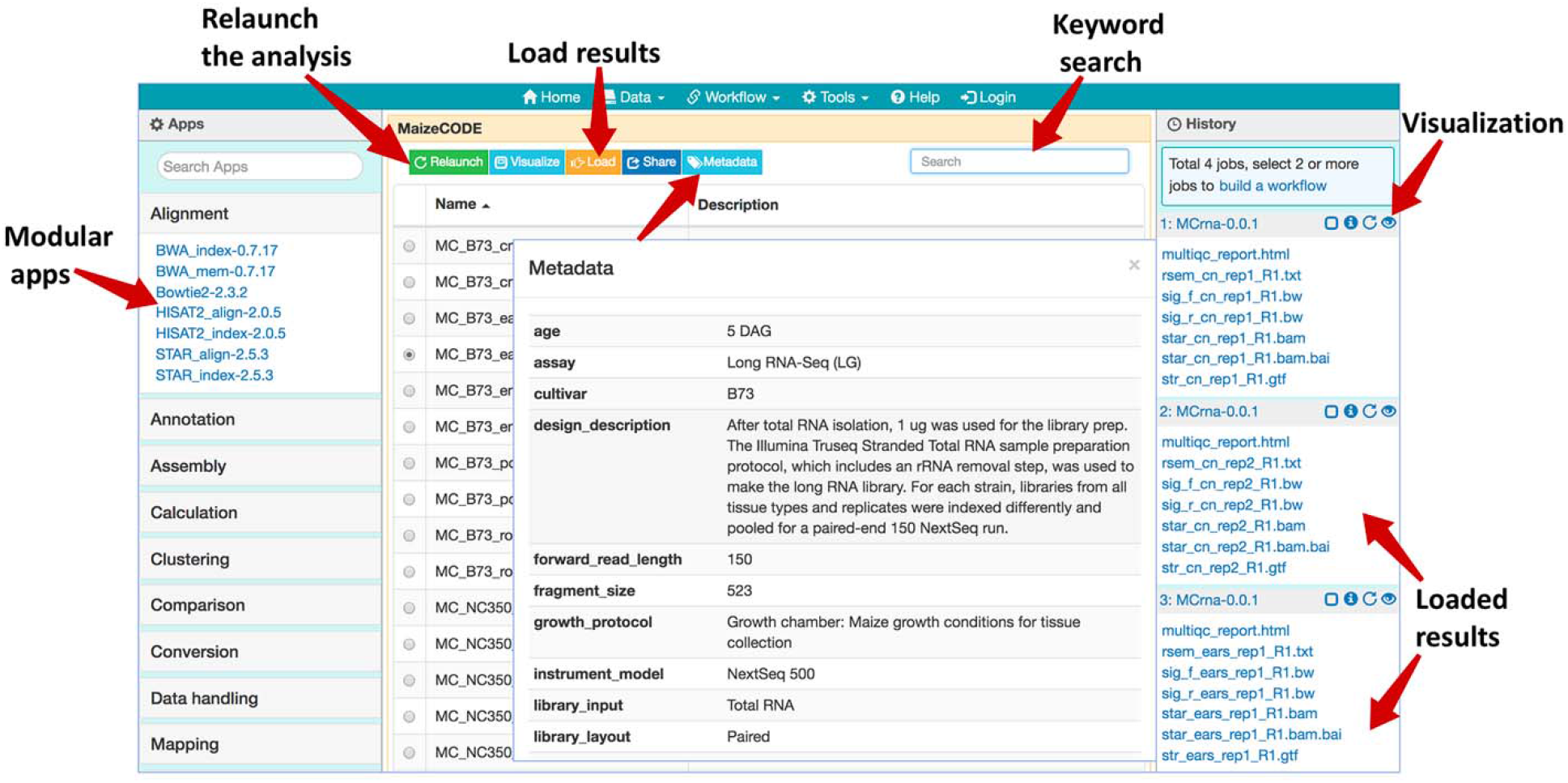
Web browser interface of the MaizeCODE data page. In the middle panel page, a list of workflows/experiments is presented. Above the list, several action buttons are available: ‘Relaunch’ the analysis, ‘Visualize’ the graphic diagram of the workflow (with URLs for the raw sequence files from the input file node), ‘Load’ the results to the History panel, ‘Share’ the analysis with others, and display the experimental ‘Metadata’. User can perform a keyword search for a specific dataset (e.g., B73 ears RNA-Seq). In the right panel, SciApps displays the history of the selected datasets; the visualization (eye) icon opens a panel where users can generate links to visualize the results in a web browser (e.g., a QC report) or genome browser (e.g., alignments or signal tracks). The left panel shows a list of modular apps that can be launched to perform a variety of downstream analyses with the loaded results.

### Accessing the MaizeCODE data as Genome Browser tracks

Once the analysis is completed, genome browser tracks are automatically generated given the workflow ID by calling the SciApps API for an integrated version of JBrowse. The browser tracks can be accessed under the ‘Tools’ menu within the top navigation bar. As shown in **Figure 2**, tracks are organized by genome, tissue, replicate, and assay. Checking the box next to each track will load it into the browser. The SciApps workflow ID is embedded, so clicking on a track brings up the workflow ‘Relaunch’ interface, which can be used to reproduce the track signal if needed. In this interface, the user can also check the parameters used for the analysis, as well as additional results in the History panel. At the bottom of the interface, a diagram button visualizes the workflow diagram, and a metadata button displays the experimental metadata associated with the workflow. From the results, user can also generate additional browser track links through the visualization (eye) icon. For example, this can be used to verify the signal track with the alignment files (in the BAM format). As mentioned earlier, the results can also be used to perform a downstream analysis on the same interface. Finally, the browser tracks are available as a JSON file for integration into other platforms (e.g., the JSON file for B73 is available at https://data.sciapps.org/view2/data2/B73/v4/apollo_data/trackList.json).

**Figure 2.**
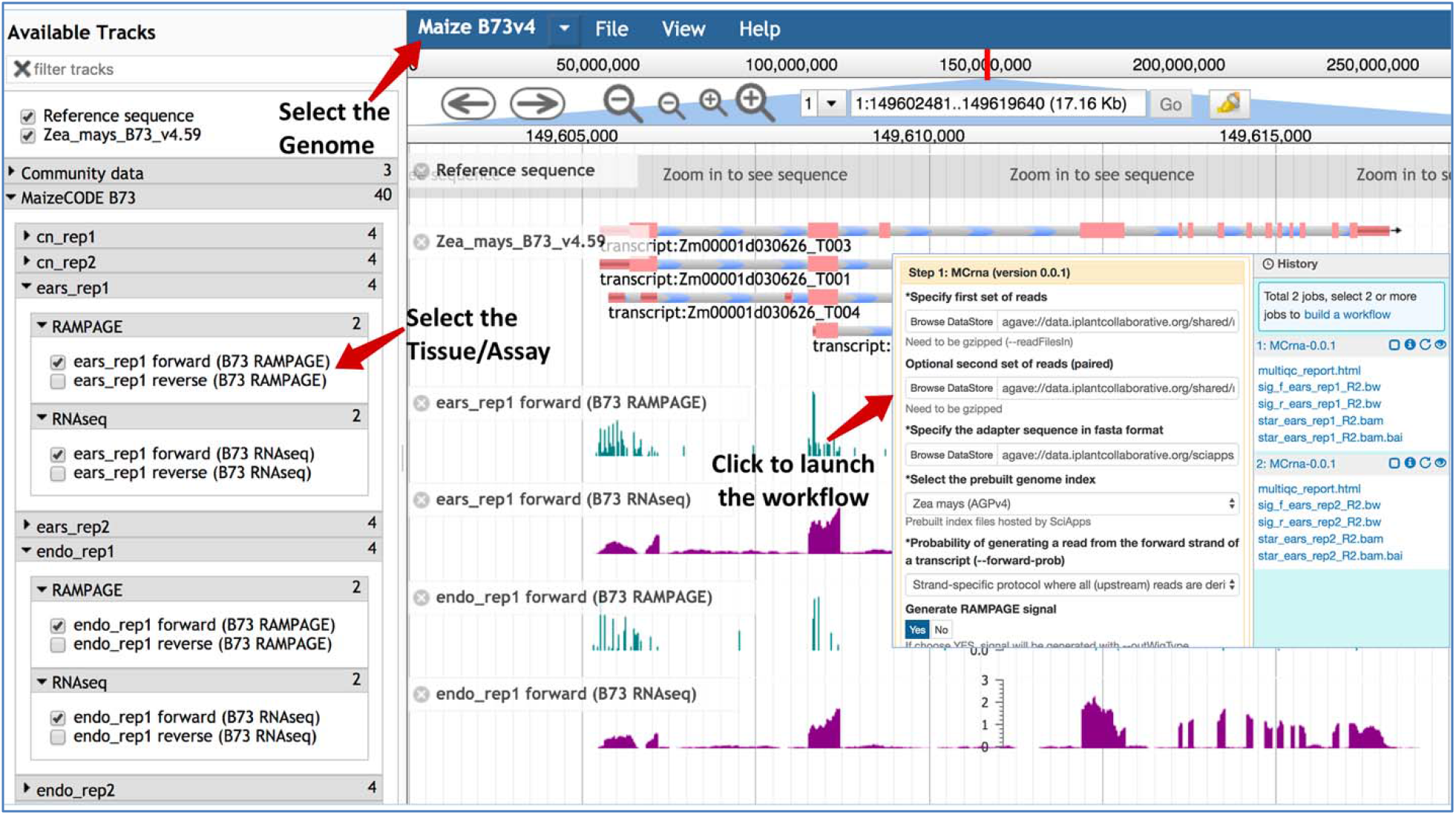
Genome browser tracks for the MaizeCODE data. JBrowse is used to hold the MaizeCODE signal tracks, which are organized in the following order: genome, tissue, replicate, and assay. Clicking on each track brings up the workflow ‘Relaunch’ interface.

Users can also locate a gene on the JBrowse by pasting the gene ID (e.g. Zm00001d02723) into the address field, as shown in Figure S3.

### Accessing the raw reads on CyVerse Data Store

The raw sequence data is deposited into the CyVerse Data Store via iCommands (https://docs.irods.org/4.2.1/icommands/user/), with metadata attached before submission to the NCBI SRA. From there, users can access the raw data in several ways. Within SciApps, the input file node of the graphic diagram for a workflow/experiment is linked to the raw sequence file. Clicking on the input node will open the CyVerse Data Common landing page in a web browser. The metadata attached to the raw sequence file is also displayed on the same page. The user can further navigate through all released raw data from the landing page (http://datacommons.cyverse.org/browse/iplant/home/shared/maizecode/released/); the SciApps workflow ID is attached as metadata to the raw data files if it has been processed. The user can use the ID to load the workflow on the SciApps portal. For batch downloading of raw sequence files through the GUI or the command line, we recommend CyberDuck (https://cyberduck.io/) or iCommands, respectively.

### Analysis with reproducible workflows

As previously described (Wang et al., 2018), bioinformatics applications (or apps) are integrated into SciApps as modular components that can be chained with other apps into an automated workflow. Individual apps are built with Singularity images (Kurtzer et al., 2017) from BioConda recipes (Grüning et al., 2018) or directly from Dockerfiles to ensure reproducibility across different cloud resources. To support the analysis of MaizeCODE data, over 20 software tools are integrated. **Figure 3** shows two publicly accessible workflows for differential expression analysis and cytosine methylation analysis, building on the popular STAR (Dobin et al., 2013)/RSEM (Li and Dewey, 2011)/StringTie (Pertea et al., 2015)/ Ballgown(Frazee et al., 2015) and Bismark (Krueger and Andrews, 2011) pipelines, respectively. These workflows can be constructed either with the SciApps GUI or through the API. The user can retrieve the inputs, metadata, results, and provenance of the software used in the analysis with a unique workflow ID. The interactive graph, along with the platform guide (https://cyverse-sciapps-guide.readthedocs-hosted.com/en/latest/index.html), helps users to understand how multiple apps are used together to analyze a specific assay. For MaizeCODE data, the graph is also helpful for visually inspecting the input–output relationship. Additionally, the user can check the input data (through the input file node) and relaunch each individual step of the analysis, or even the entire analysis, via the web interface or API.

**Figure 3.**
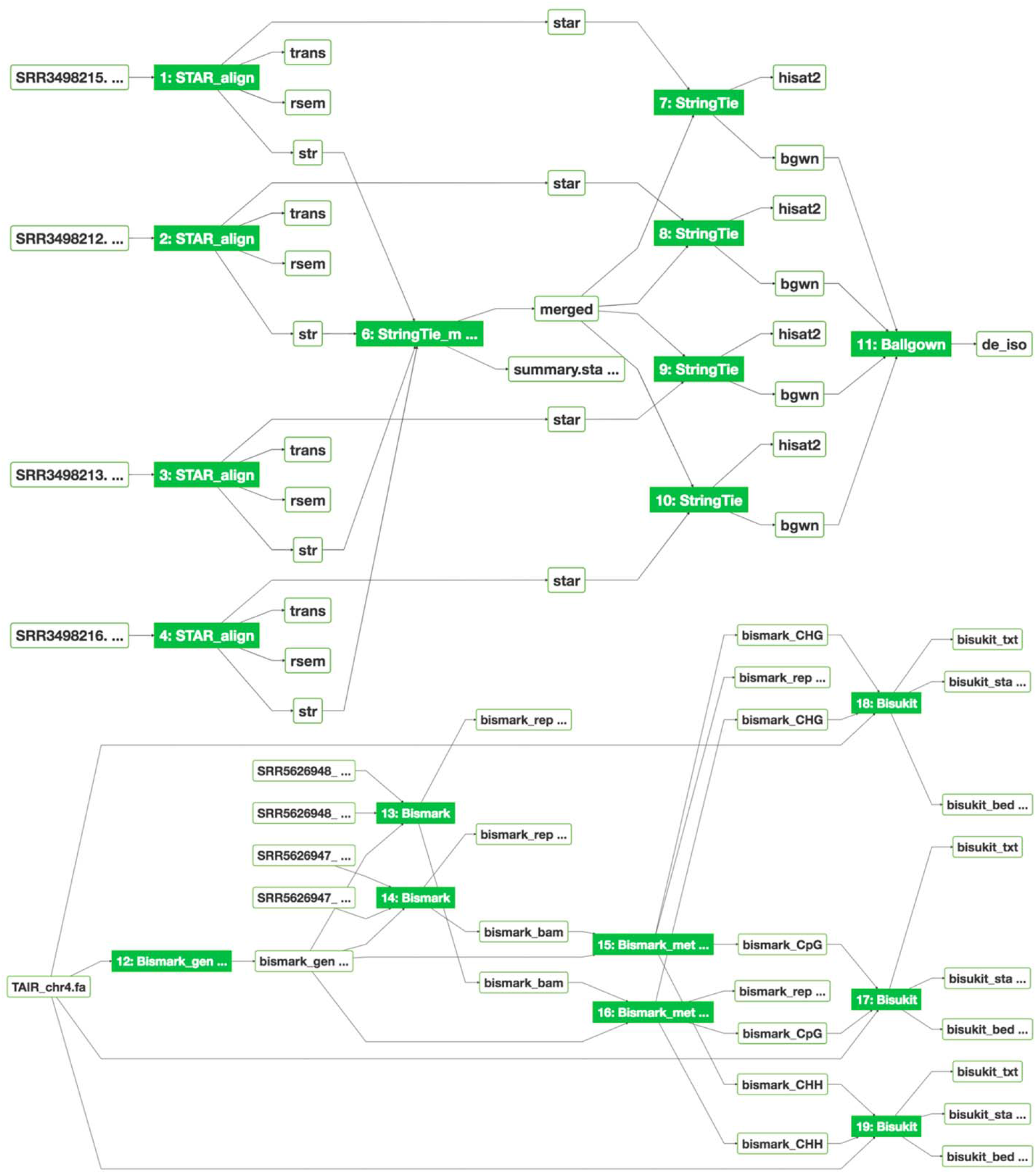
Graphical workflow diagrams for differential expression analysis (top) and MethylC-seq analysis (bottom). The interactive graph demonstrates the relationships among input–output files, displays provenance of the software tools, and provides real-time job status updates of new analyses via the color of the app node (green: completed; blue: running; yellow: pending; red: failed).

## RESULTS AND DISCUSSION

A large variety of software is needed to process the MaizeCODE data. For each experiment (consisting of two replicates), a workflow with a unique ID is provided via the SciApps platform. One major goal of SciApps is to empower anyone in the community to easily repeat an entire analysis, or use a workflow with alternative parameters for each step if so desired. A second major goal is to empower community members to process and combine their own comparable data sets if they are generated with similar protocols used by the MaizeCode project, available at https://datacommons.cyverse.org/browse/iplant/home/shared/maizecode/released/protocols. In the following sections, we describe how RNA-seq data is processed, how the results can be visualized, and how the primary analysis results can be used for differential expression analysis.

### Processing the RNA-seq data

Besides the UCSC genome analysis tool bedGraphToBigWig (for format conversion and generating browser track signals), the major software used in MaizeCODE RNA-seq data analysis are bbduk (https://jgi.doe.gov/data-and-tools/bbtools/bb-tools-user-guide/bbduk-guide/) for trimming low quality reads and adapter sequences, FastQC and MultiQC (Ewels et al., 2016) for visually checking read quality, STAR (Dobin et al., 2013) for alignment, RSEM (Li and Dewey, 2011) for quantifying gene expression, and StringTie (Pertea et al., 2015) for transcriptome assembly. All tools are integrated into SciApps individually as separate apps, and also combined as a single app, MCrna, for rapid batch processing of RNA-seq data without requiring intermediate results to be transferred between the TACC and CyVerse cloud. **Figure 4** shows the relationship among these analysis tools within the MCrna app, which is used to process both replicates of an experiment in parallel.

**Figure 4.**
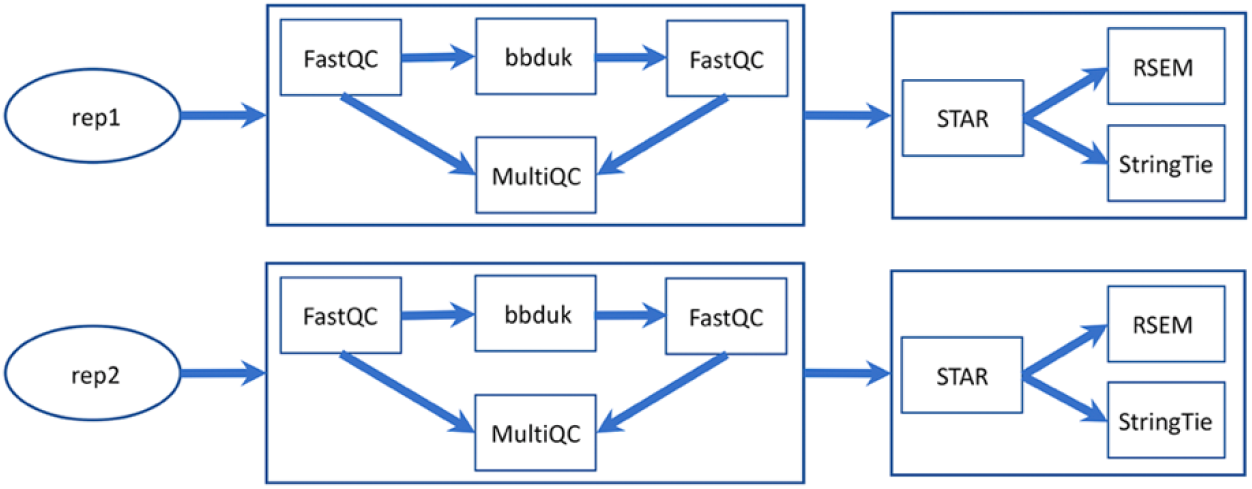
MaizeCODE MCrna app for processing RNA-seq data from two replicates. The MCrna app wraps six tools, FastQC, bbduk, MultiQC, STAR, RSEM, and StringTie, together for QC and quantification of each replicate.

For each replicate, the MultiQC software outputs a quality report for the sequence data, before and after trimming, in an interactive HTML format. This report can be accessed via the visualization (eye) icon in the History panel (next to each loaded replicate, as shown in **Figure 1**). As with the HTML format, text, image, and other web browser–compatible files can be visualized by clicking the icon. For files that can be displayed on a Genome Browser (e.g., BAM, bigwig, etc), the user can also generate browser–ready links by clicking the same icon. These links address the cloud storage system from the CyVerse project, so they can be displayed on Genome Browsers hosted by different portals. If the user clicks on the output file name (from the History panel), they will be directed to the CyVerse Data Commons landing page, where they can preview or download the results. For files over a few GBs, we recommend that the user download their data using either iCommands or CyberDuck, using the CyVerse Data Store path available in the file URL.

### Automated differential expression analysis

One of the key advantages of distributing the MaizeCODE data through SciApps is that it facilitates downstream analysis. In this section, we will show how differential expression analysis can be performed, on either the gene or isoform levels, using the primary analysis results discussed in the last section.

As shown in **Figure 5**, after loading the results for two experiments from the MaizeCODE data page (**Figure 1**), one for ear tissue and the other for the root tissue, we can launch the RSEM_de app from the ‘Comparison’ category (or through searching the app panel). For the analysis, users drag and drop output files with names starting with ‘rsem_’ into the input field for both replicates of each sample. The analysis job can then be submitted to the cloud for running, and the results (i.e., the differentially expressed genes) will be available within a few minutes. Note that the app is flexible in handling different numbers of replicates per sample. Additional input fields can be added using the ‘+ Insert’ button. For the MaizeCODE project, most data sets are generated with two replicates.

**Figure 5.**
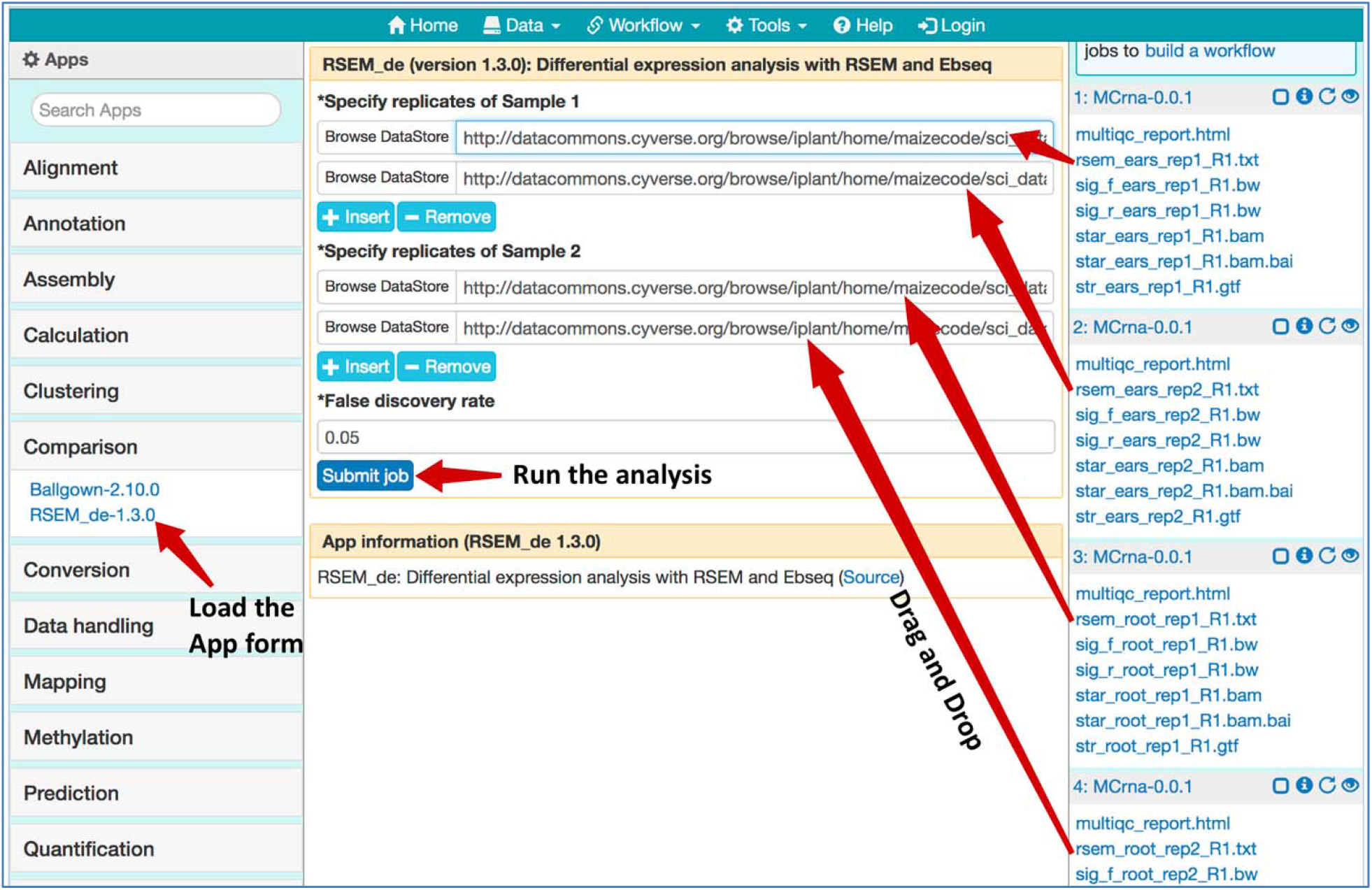
Using the RSEM_de app for gene-level differential expression analysis. Job histories are “Loaded’ from the MaizeCODE data page (https://www.sciapps.org/data/MaizeCODE); Clicking the RSEM_de-1.3.0 from the left App panel brings up the app form in the main panel; Dragging and dropping the gene quantification result files starting with “rsem_” into the input field, then clicking the “Submit job” button to run the differential expression analysis.

Users can check the results through the History panel or the list of jobs (under the ‘Workflow’ tab from the top menu). Users can also select jobs from the History panel and save them as new workflows to organize the analysis and/or share it with others. Given that the XSEDE/TACC cloud is a shared resource, and jobs may be queued for several minutes to several hours, we have also established a local cluster (Wang et al., 2015) to quickly process small jobs requiring less than an hour to complete. Powered by the Agave API (Dooley et al., 2012), SciApps treats both the XSEDE/TACC cloud and the local cluster as virtual execution systems, allowing a scientific app to be configured to run on either cloud. As such, by using the CyVerse Data Store as a central storage hub, SciApps workflows can be seamlessly executed on a mixture of XSEDE/TACC cloud and local clusters for efficient yet scalable processing and consumption of MaizeCODE data.

To perform differential expression on the isoform level, users can launch the workflow diagram shown in **Figure 3**. From the diagram and the relaunched app forms, the StringTie_Merge app is used to merge assembled transcripts from all replicates to generate a new annotation file. The annotation file is then passed again to the StringTie app to compute gene expression quantification results, which are then input to the Ballgown app (Frazee et al., 2015) to compute differentially expressed isoforms. Again, the user can drag and drop the alignment files and assembled transcripts from the MaizeCODE primary analysis results without repeating the time-consuming alignment and quantification steps. Given that all results are accessible through web URLs, users can also retrieve the data directly to their local server for further interactive analysis. For example, SciApps users can inspect the quantification results of each pair of replicates to confirm that they are consistent, and if they are, proceed with the analysis of differentially expressed genes.

## CONCLUSION AND FUTURE WORK

SciApps is a cloud-based platform that provides a data management infrastructure for the MaizeCODE data. The platform supports the management of experimental metadata and computational provenance; provides a collection of analysis apps covering analysis of multi-omics data sets; and provides both web browser and API access to data, results, metadata, and computational provenance of software tools through a unique workflow ID. Genome browser tracks are also provided to enable visualization of the results using an integrated version of JBrowse.

SciApps has been designed to integrate cloud resources for scaling and long-term stability. Currently, SciApps has been integrated with the NSF-funded XSEDE/TACC cloud and the CyVerse Data Store to provide academic users with cost-free data storage and computing services. Both resources are integrated as virtual systems via the Agave API, which also supports the integration with commercial cloud platforms like Amazon EC2/S3 and Microsoft Azure. Therefore, SciApps can be scaled for large-scale data management and analysis if needed. Additionally, SciApps can also seamlessly integrate local data and computing resources to complement cloud resources. The successful utilization of our local system suggests that SciApps can facilitate collaborative projects across different institutes for joint data production, analysis, and management with multiple local systems, thereby avoiding high financial costs.

To process the data sets that are continually being generated by the MaizeCODE project, several new analysis tools have been integrated into SciApps. Future goals include developing new analysis workflows, supporting sophisticated queries against metadata, reanalyzing and distributing published sequencing data sets from raw data, and conducting training and community outreach.

## IMPLEMENTATION

As previously described, the major components of the SciApps analysis portal include a web browser user interface, a MySQL database, a workflow engine, and a newly developed web API. The web API is written in Perl to complement the web browser user interface, especially for batch processing and metadata handling. Analysis jobs are submitted to the cloud, and in addition all released raw data, processed results, metadata, and workflows, are made publicly accessible through both the GUI and API with no authentication needed.

To support MaizeCODE data management and enhance SciApps functionalities, several searchable pages are added, including the user workflow page, user job page, and MaizeCODE data page. On these pages, users can relaunch the workflow/job, visualize the workflow diagram, load the job history and results into the History panel, share the workflow with others, and check the metadata associated with the workflow. The RESTful API is designed to facilitate batch processing of the MaizeCODE data through template workflows. More details and other updates to the SciApps platform are described in the Supplementary file section 1 (page 2). The overall cycle of MaizeCODE data management and processing is described in Figure S2.

## AUTHOR CONTRIBUTIONS

LW and ZL designed, implemented, and tested the software. MD and LW managed metadata and submission of raw data to NCBI SRA. LW and PVB designed and maintained the local system consisting of a web server, a data server, and several computing servers. LW, ZL, and XW developed the scientific workflows and participated in testing through the web interface. AD and TG helped in developing the workflow. XW and LW processed the data with the automated workflows. MR, JD, XX, COR, KB, DJ, RM, SG, and WM generated the raw data. CG developed the Perl code for interacting with Agave API. CFM, CG, and DM managed the MaizeCODE website. LW, DW, MS, and TG wrote the manuscript. All the authors read and approved the final manuscript.

## AVAILABILITY AND REQUIREMENTS

Project name: SciApps

Project home page: https://sciapps.org/

Source code repository: https://github.com/warelab/sciapps

Tutorial: https://cyverse-sciapps-guide.readthedocs-hosted.com/en/latest/maizecode.html

Software Design Document: https://github.com/warelab/sciapps/blob/master/doc/SDD.pdf

MaizeCODE data page: https://www.sciapps.org/data/MaizeCODE

Operating system(s): Platform independent

Programming language: JavaScript, Perl

License: MIT

Any restrictions to use the data: Toronto Agreement

## FUNDING

This work is supported by NSF grant IOS 1445025 for MaizeCODE. SciApps has also been supported by NSF grant DBI-1265383, and USDA-ARS (1907-21000-030-00D).

## Notes

#### Summary of Updates

The title is changed and the first figure is removed.

https://www.sciapps.org

